# Physical contacts between sparse biofilms promote plasmid transfer and generate functional novelty

**DOI:** 10.1101/2023.02.01.526699

**Authors:** Josep Ramoneda, Yinyin Ma, Julian Schmidt, Michael Manhart, Daniel C. Angst, David R. Johnson

## Abstract

The horizontal transfer of plasmids is an important driver of microbial evolution, such as conferring antibiotic resistance (AR) to new genotypes. In biofilms, the abundance of cell-cell contacts promotes the frequent transfer of plasmids and their associated genes. In this study, we expand our knowledge about AR-encoding plasmids by investigating their transfer between discrete biofilms as the biofilms grow and physically collide with each other. Using an experimental system consisting of two fluorescently labelled *Pseudomonas stutzeri* strains and an *Escherichia coli* strain, we show that biofilm collisions promote plasmid transfer along the collision boundaries. The extent of plasmid transfer depends on the plasmid loss probability, the plasmid transfer probability, and the relative growth rates of plasmid-free and plasmid-carrying cells. We further show that the proliferation of plasmids after biofilm collision depends on the spatial positionings of plasmid-carrying cells along the collision boundary, thus establishing a link between the large-scale spatial distribution of discrete biofilms and the small-scale spatial arrangement of cells within individual biofilms. Our study reveals that plasmid transfer during biofilm collisions is determined by spatial factors operating at different organizational levels and length scales, expanding our understanding of the fate of plasmid-encoded traits in microbial communities.

## Introduction

Microbial communities growing across surfaces are pervasive on our planet [1], drive important biogeochemical cycles [2, 3], and affect human health and disease [4–6]. When embedded in a matrix of extracellular polymeric substances, these so-called biofilms are involved in a myriad of biotechnological applications such as water decontamination and biofuel production [7, 8], but also cause persistent infections in animal tissues and contaminate medical devices [9]. Within biofilms, the close spatial proximities of individual cells drive multiple processes, among which is the horizontal transfer of plasmids and their associated genes (i.e., circular pieces of DNA that often contain functionally important genes such as antibiotic resistance [AR]) [10]. The processes of plasmid loss (errors in segregation control upon cell division) and horizontal transfer (conjugation) are the main determinants of plasmid fate and proliferation within actively growing biofilms [11]. AR-encoding plasmids can cause persistent AR bacterial populations in human and environmental microbiomes, posing a serious threat to global health [12, 13]. As biofilms are hotspots for plasmid transfer [14], it is important to understand how the spatial features of biofilms drive the spread of AR-encoding plasmids.

Despite the recognition of biofilms as hotspots for plasmid transfer, the majority of transfer is postulated to occur along the outer edges of the biofilm matrix [11]. This is because plasmids generally only invade into and subsequently proliferate within metabolically active cells, which are usually those cells lying at the outer edges of the biofilm where unoccupied space and nutrients are plentiful [15]. This expectation has been confirmed in natural systems such as the mouse gut [16], where transfer occurs only at the edges of the mucus layer covering epithelial cells. Recent studies on bacterial communities growing across nutrient-rich agar surfaces, however, show that plasmid transfer is pervasive within biofilms as long as cells are actively growing (i.e., undergoing range expansion, [17, 18]). To better understand plasmid dynamics in biofilms, it is thus necessary to delineate plasmid transfer occurring within biofilms and at the biofilm boundaries, and to determine whether plasmid transfer in these scenarios is driven by the same or different mechanisms.

In addition to space and nutrient availability, the spatial arrangement of cells across a surface is another important determinant of plasmid transfer and spread [15, 17, 18]. During biofilm growth and concomitant expansion across surfaces, the component microbial populations typically spatially segregate from each other as a consequence of drift at the expansion edge [19]. This process has important effects on biofilm diversity [20, 21], stability [22], and functioning [23]. Plasmids transfer to greater extents within communities with highly spatially intermixed populations [18]. This effect is ascribed to the higher number of cell-cell contacts between plasmid-free and plasmid-carrying cells, which increases the number of possible plasmid transfer events [11, 24, 25]. Frequent disturbance is a factor that can promote plasmid invasion by causing the spatial reorganization of cells and creating new cell-cell contacts in otherwise spatially-segregated populations [15, 26]. However, in a given environment surfaces are generally not colonized by a single contiguous biofilm but are rather colonized by multiple spatially segregated biofilms (i.e., sparse biofilms), where each discrete biofilm lies adjacent to others and dynamically expands and contracts in size as a consequence of growth and death [27, 28].

Despite the implications for the dissemination of AR, there is surprisingly little information on the processes governing plasmid transfer between adjacent biofilms [11, 29, 30]. Scenarios where discrete biofilms expand and eventually collide into each other are likely common in systems such as the gut lumen [31–33] and the dental plaque [34, 35]. The processes of biofilm expansion, collision, and retraction are more prominent when periodically exposed to disturbances such as antibiotics, where antibiotic administration can drastically reduce the population sizes of sensitive individuals while also exacerbating the subsequent spread of plasmid-encoded AR during biofilm recovery by imposing a positive selection pressure [36, 37]. Understanding the mechanisms driving AR-plasmid transfer during biofilm collisions would thus fill a knowledge gap by linking the dynamics within individual biofilms to the dynamics between biofilms.

In this study, we investigated the determinants of the horizontal transfer of AR-encoding plasmids during biofilm collisions. To accomplish this, we developed a novel experimental system in which one biofilm, consisting of a pair of fluorescently labelled strains of the bacterium *Pseudomonas stutzeri* A1501 that carry a conjugative plasmid pAR145 encoding for chloramphenicol resistance, expands and eventually collides with an *Escherichia coli* biofilm. We performed biofilm collision experiments by colliding the two-strain *P. stutzeri* biofilm with the one-strain *E. coli* biofilm, which allowed us to study both intraspecific (within the two-strain biofilm) and interspecific (between the two-strain and one-strain biofilms) plasmid transfer in the most simplified manner. We aimed to 1) quantify the extent of plasmid transfer from the *P. stutzeri* biofilm (donor biofilm) to the *E. coli* biofilm (recipient biofilm) upon collision, and 2) determine how the cellular-level organizations of the colliding biofilms affect plasmid transfer and its subsequent spread. We experimentally identified the determinants of plasmid transfer to *E. coli* upon biofilm collision and the processes leading to the subsequent proliferation of these new genotypes within the recipient *E. coli* biofilm. We further examined the mechanisms by which spatial factors determine the proliferation of new genotypes after biofilm collision using an individual-based computational model.

## Results

### pAR145 dynamics prior to biofilm collision

We first quantified pAR145 dynamics within the *P. stutzeri* biofilm prior to collision with the *E. coli* biofilm. We expected the processes of pAR145 loss and transfer during expansion of the *P. stutzeri* biofilm alone to have a consequential role on the potential transfer of pAR145 to the recipient *E. coli* biofilm. We found that pAR145 was purged from the *P. stutzeri* biofilm within a 300 μm distance corresponding to the radial interval between 1700-2000 μm (Figs. 1A and 1B), which is an accumulated outcome of pAR145 loss and preferential growth of pAR145-free individuals in the absence of chloramphenicol. In our system, this window determines the space and time during which the transfer of pAR145 to the *E. coli* recipient biofilm will be maximal when colliding with an adjacent biofilm.

**Figure 1.**
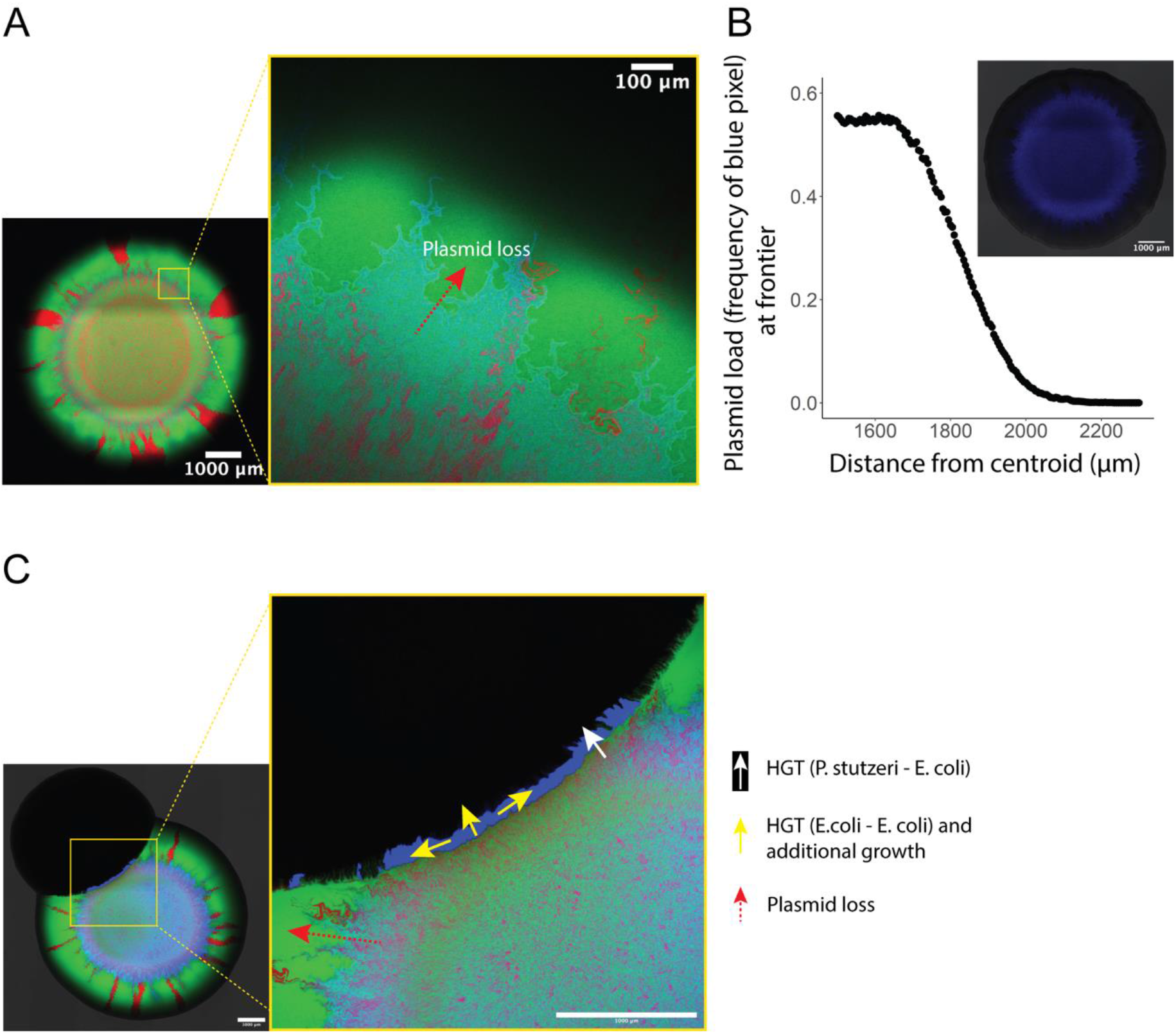
Plasmid dynamics in expanding biofilms and transfer into adjacent biofilms upon collision. **A,** Representative microscopy image of the *P. stutzeri* biofilm consisting of two isogenic strains (one expresses red fluorescent protein while the other expresses green fluorescent protein) that expand together. Initially, the pAR145 donor strain carried pAR145 (purple) while the potential recipient did not (green). Note the rapid demixing of the two genotypes and the formation of discrete sectors. pAR145 transfer can occur between the two *P*. *stutzeri* strains, upon which the green strain will turn cyan. The red dashed arrow indicates a pAR145 loss event and subsequent proliferation of pAR145-free cells. **B,** Quantification of pAR145 abundance (blue fluorescence signal) during range expansion of the *P. stutzeri* donor biofilm shown in panel A. **C,** The left panel is a representative microscopy image after physical collision between the *P. stutzeri* donor and *E. coli* recipient biofilms. Note the formation of a blue patch located at the collision boundary, which corresponds to *E. coli* cells that acquired pAR145 via transfer from pAR145-carrying *P. stutzeri* cells. The exposure of the blue channel has been increased to better visualize the boundaries between plasmid-carrying and -free cells. The right panel is a magnified image of the collision boundary. White arrow: successful pAR145 transfer from the *P. stutzeri* donor biofilm to the *E. coli* recipient biofilm. Yellow arrows: pAR145 transfer and proliferation within the *E. coli* recipient biofilm. Red arrow: pAR145 loss within the *P. stutzeri* donor biofilm.

### pAR145 transfer between colliding biofilms

We next verified that physical collisions between the *P. stutzeri* donor biofilm and the *E. coli* recipient biofilm can promote pAR145 transfer into the *E. coli* biofilm. To accomplish this, we inoculated a 1 μL droplet of the donor *P. stutzeri* consortium at ca. 3.0 mm from the recipient *E. coli* biofilm. After 96h of incubation, the *P. stutzeri* donor and *E. coli* recipient biofilms had physically collided, and a new genotype formed at the collision boundary (Fig. 1C). This new genotype is the initially non-fluorescent *E. coli* that obtained pAR145 and expresses cyan fluorescent protein upon contact with the pAR145-carrying *P. stutzeri* biofilm. A closer evaluation of the collision boundary revealed that the newly formed genotype homogenously extended across an approximately 2 mm long boundary and protruded ca. 80 μm into the *E. coli* biofilm (Fig. 1C).

### pAR145 loss and initial biofilm positioning determine pAR145 transfer during biofilm collisions

After verifying that pAR145 can transfer between biofilms upon their physical collision, we next investigated the main determinants of this process. We hypothesized that the initial distance between the *P. stutzeri* donor and *E. coli* recipient biofilms would determine the extent of pAR145 transfer, where larger distances increase the time for pAR145 loss prior to biofilm collision and thus reduce pAR145 transfer. To test this hypothesis, we inoculated the *P. stutzeri* donor and *E. coli* recipient biofilms at precisely defined distances from each other. We found that collisions between biofilms initially inoculated at closer distances formed larger amounts of new *E. coli* transconjugants upon collision (ANOVA *F*_1, 28_ = 28.95, *P* = 9.8 x 10^-6^, n = 5) (Figs. 2A and 2B). The spatial range in which new *E. coli* transconjugants were formed were initial distances of 2.7-3.4 mm, although the amount new *E. coli* transconjugants decreased monotonically within this window (Spearman’s rank correlation coefficient = −0.793; *P* = 8.6 x 10^-5^) (Fig. 2B).

**Figure 2.**
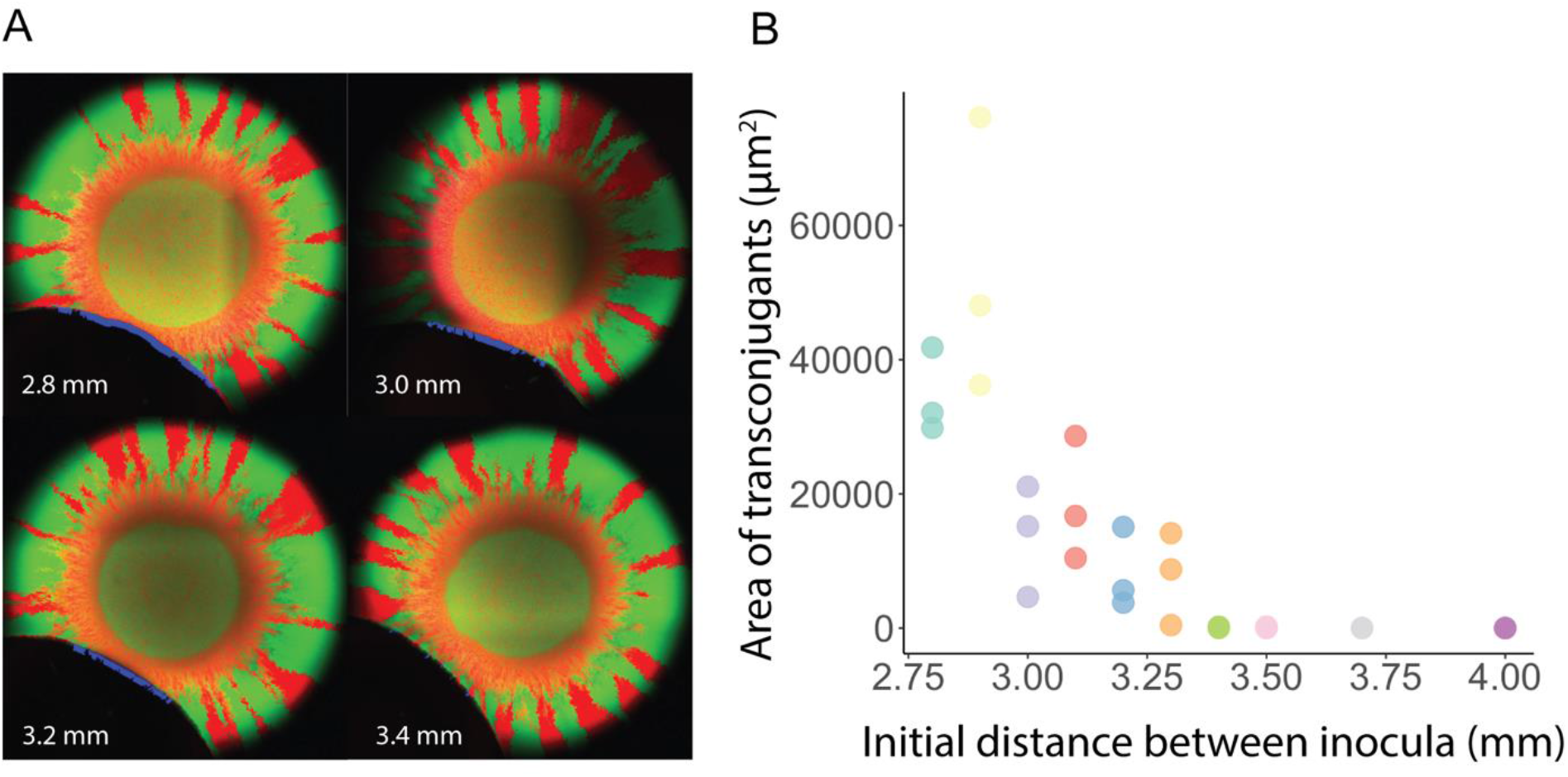
Initial distance between biofilms determines the extent of pAR145 transfer upon biofilm collision. **A,** Representative microscopy images for experimental collisions between *P. stutzeri* donor and *E. coli* recipient biofilms. Numbers indicate the spatial distances between the initial inocula. pAR145 transfer occurs at the collision boundaries and generates blue patches consisting of *E. coli* transconjugants carrying pAR145. Note that due to overexposure of the green and red channels, mixed populations of *P. stutzeri* do not display visible blue signals even when carrying pAR145. In contrast, *E. coli* does not express green or red fluorescent protein and appears blue when carrying pAR145. **B,** Quantification of the absolute area of *E. coli* transconjugants as a function of the distance between the initial inocula. Each datapoint is for an independent biological replicate (n = 3) at the specified initial distance (note that some datapoints are overlapping and thus appear as one). As the initial distance between the inocula increases, the area of newly created transconjugants declines.

### What is the relative contribution of plasmid transfer versus cell proliferation on plasmid spread upon collision?

We next sought to recapitulate our findings, quantify the relative contributions of plasmid transfer (HGT) and cell proliferation to the total number of transconjugants, and understand the factors limiting the extent of HGT using an individual-based computational model. We found that the biophysical modelling framework defined in CellModeller [38] (see Materials and Methods) successfully captured the formation of transconjugants within the recipient biofilm upon biofilm collisions (Fig. 3A). Similar to the experimental results, the distance between the initial inocula had a strong impact on plasmid transfer upon collision between the simulated biofilms (Fig. 3A). Both the total number of new transconjugants (ANOVA *F*_1, 28_ = 143.6, *P* = 1.5 x 10^-12^, n = 5) (Fig. 3B) and the number of transfer events from the donor to the recipient biofilm (ANOVA *F*_1, 28_ = 93.9, *P* = 1.9 x 10^-10^, n = 5) (Fig. 3C) increased significantly at closer distances between the initial inocula.

**Figure 3.**
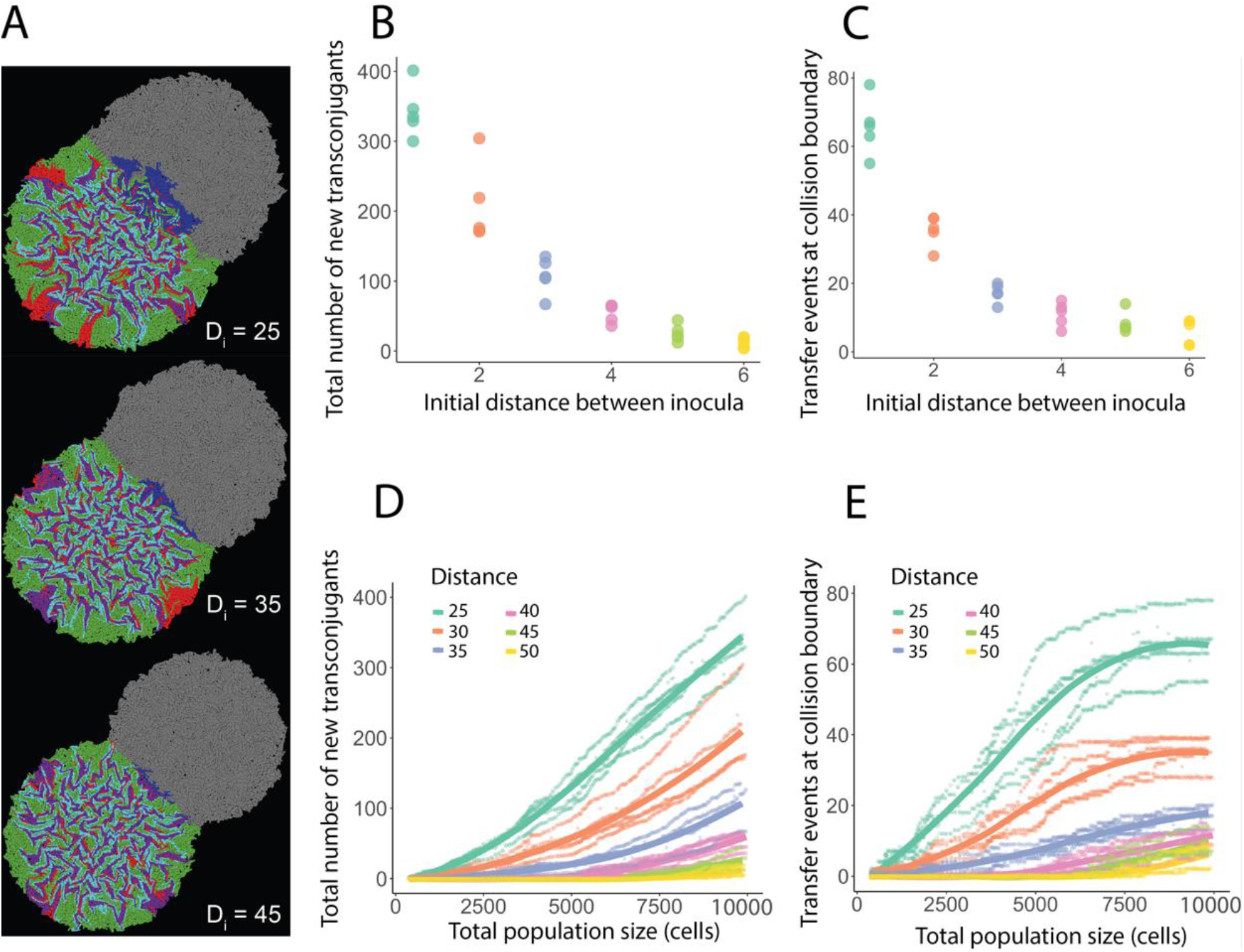
Simulated collisions leading to plasmid transfer and spread between a donor and recipient biofilms expanding at different initial distances. **A,** Representative simulations where the initial inoculum for the donor biofilm consists of a 1:1 mixture of plasmid-carrying (purple) and plasmid-free (green) cells. Green cells have a 17% growth advantage over purple cells, which corresponds to the 17% growth rate cost for carrying pAR145 in the experiments. At each cell division there is a probability of 0.0005 that plasmid carriers will lose the plasmid due to errors in segregation control. The initial inoculum for the recipient biofilm consists of a single population of plasmid-free cells (grey) which can receive the plasmid upon collision with the donor biofilm. Once physical contact occurs, there is a probability of 0.005 at each time step that the recipient will receive the plasmid and become a transconjugant. If grey cells receive the plasmid, they will become blue (interspecific transfer). Green cells have an equal probability to acquire plasmids and turn into cyan transconjugants (intraspecific transfer). White numbers D_i_ at the bottom right of the simulation images are the distances between the initial inocula. The initial number of grey cells inoculated is the same as the initial total number of green + purple cells. Simulations were performed for 300 time steps until reaching a final population size of 10000 cells. **B,** Total number of blue transconjugants at the end of the simulations. Each datapoint is for an independent biological replicate (n = 5) (note that some datapoints are overlapping and thus appear as one). Different colors indicate different distances between the initial inocula. **C,** Number of transfer events at the collision boundary quantified at end of the simulations. Transfer events refer to interspecific transfer between the donor and recipient biofilms. **D**, Total number of blue transconjugants upon biofilm collision. **E**, Number of interspecific transfer events at the collision boundary as a function of the total population size. For both panels, each data point is for an independent simulation (n=5) and the solid lines are the running averages. Different colors are for different initial distances between the initial inocula.

Using the modelling framework we tracked the formation of new transconjugants during biofilm development. We first investigated how the total number of new transconjugants changed during biofilm development and found a steady increase in the accumulated number of new transconjugants throughout the simulations (Fig. 3D). This is largely because plasmid transfer continues to occur between transconjugants and their ancestral recipients (intraspecific transfer). Therefore, we next evaluated the number of interspecific transfer events that accumulate along the collision boundary and found that these events increased significantly from the beginning of biofilm development but then flattened out as the simulations continued (Fig. 3E). The cause of this saturation is that successful transfer between two expanding biofilms has largely saturated at the collision boundary (i.e., there are no longer any remaining cell-cell contacts between the donor and recipient populations for which plasmid transfer has not occurred), and thus new transconjugants can no longer be generated.

### Spatial intermixing determines plasmid spread after biofilm collision

In addition to the absolute number of plasmid donors along the collision boundary, we hypothesized that spatial intermixing of different populations along the collision boundary is also an important determinant of plasmid spread after biofilm collision. As the distance between the initial inocula increases, we expected plasmid-carrying and plasmid-free cells to become increasingly spatially segregated due to longer expansion times (as reported in [17, 18]), and lead to less efficient plasmid transfer both between biofilms and within the recipient biofilm upon collision. We tested two potential effects associated with the spatial intermixing of plasmid-carrying and plasmid-free cells that can modulate the extent of plasmid spread into an adjacent biofilm. First, we considered the effects of spatial intermixing of plasmid-carrying and -free cells in the donor biofilm on the total plasmid load at the collision boundary. Spatial intermixing determines the total number of plasmids that can be potentially transferred upon collision with an adjacent plasmid-free biofilm by modulating the efficiency of HGT (Fig. 4AB). To test this effect, we allowed for HGT to happen within the donor biofilm and from donor to recipient, but not within the recipient biofilm. Second, we considered the effects of spatial intermixing of plasmid-carrying and -free populations in the donor biofilm at the collision boundary on the intermixing of new transconjugants (blue cells) and plasmid-free cells (grey cells) in the recipient biofilm. Spatial features of the donor biofilm are acquired by the recipient biofilm upon collision and drive plasmid spread within the recipient biofilm by again modulating the efficiency of HGT (Fig. 4AC). To test this effect, we allowed for HGT to happen between the donor biofilm and the recipient biofilm, and within the recipient biofilm, but not within the donor biofilm.

**Figure 4.**
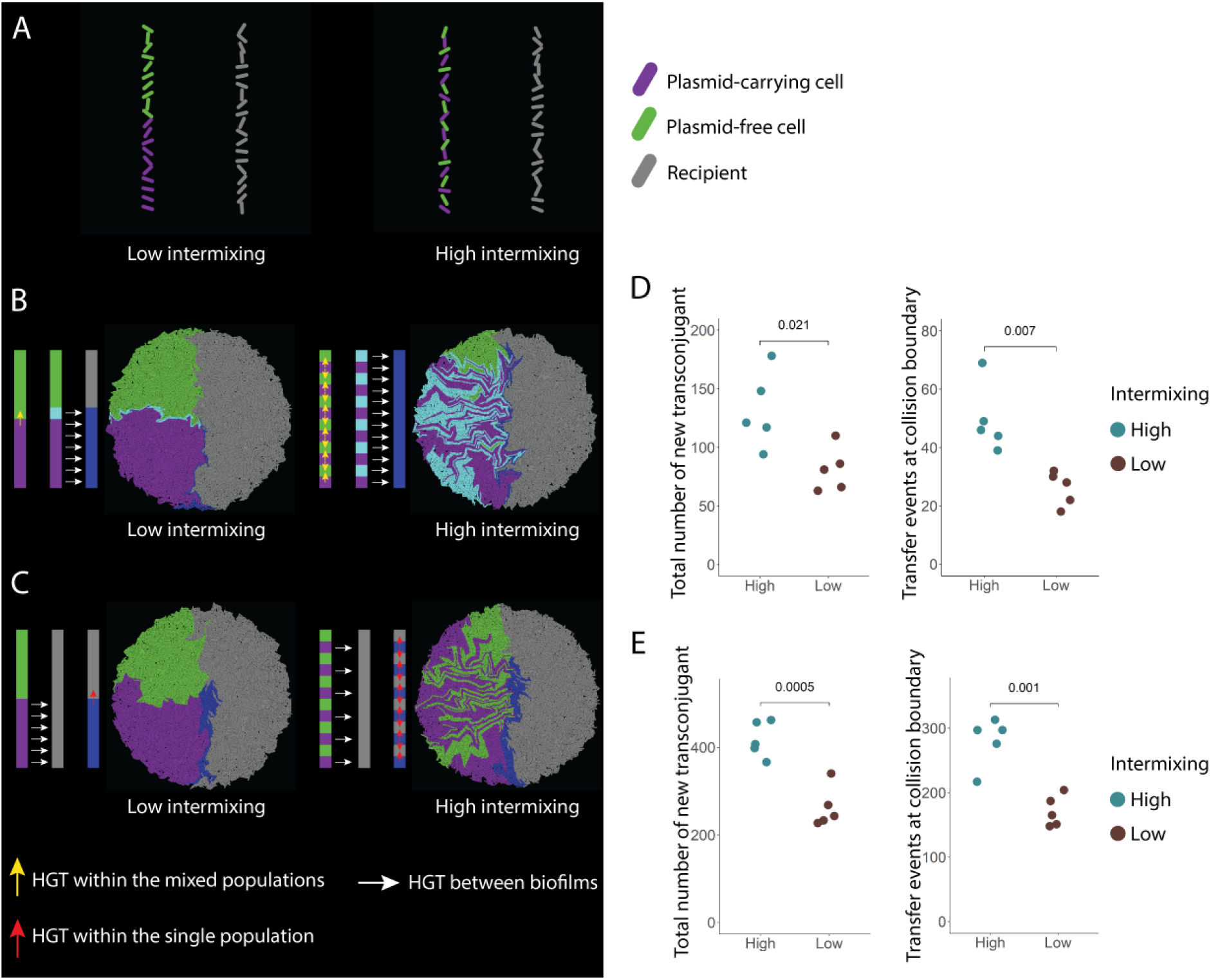
Effects of the spatial intermixing of plasmid carriers and plasmid-free individuals in the donor biofilm on subsequent plasmid spread in the recipient biofilm. **A,** Initial positioning of cells along a simulated collision boundary. Plasmid-carrying cells (purple) and plasmid-free cells (green) within the donor biofilm are positioned either in two discrete patches or are highly intermixed. Grey cells are the potential recipients within the recipient biofilm. Cells are all randomly rotationally oriented along the x-y plane. **B,** Schematic figures of the potential effects on the left and the simulations on the right. Blue cells indicate the newly created transconjugants in the recipient biofilm. Simulations tested the effects of intermixing within the donor biofilm on plasmid load at the collision boundary and on subsequent plasmid spread within the recipient biofilm. To simulate these effects, we only enabled plasmid transfer within the donor biofilm (mixed populations of purple and green) indicated by vertical yellow arrows, and between biofilms indicated by horizontal white arrows. **C,** Simulations testing the effects of intermixing within the donor biofilm on consequent intermixing within the recipient biofilm and further plasmid spread within the recipient biofilm. Blue cells indicate the newly created transconjugants in the recipient biofilm. To simulate these effects, we only enabled plasmid transfer between biofilms indicated by the horizontal white arrows, and within the recipient biofilm (grey cells) indicated by the vertical red arrows. **D, E,** Quantification of the total number of new transconjugants formed in the recipient biofilm (blue) and the total number of plasmid transfer events that occurred at the collision boundary under the simulation condition shown in **B, C**, respectively. Simulation images correspond to the final time step at 7000 total cells. P-values from Welch two-sample t-tests are shown on top of panels D and E.

We used our individual-based modelling framework to simulate the collision boundary as two “cell walls”, where one cell wall contains mixed populations consisting of plasmid-carrying cells (purple cells) and plasmid-free cells (green cells), and the other cell wall consists of one single plasmid-free population (grey cells) (Fig. 4A). We varied the initial intermixing of the mixed populations by either placing two cell types (purple and green) into two discrete patches (low intermixing), or by sequentially placing one cell type next to the other (high intermixing) (Fig. 4A). Compared to initiating cell inocula as in Fig. 3, initiating two cell walls allowed us to precisely control the spatial intermixing upon collision and investigate mechanisms occurring right at the collision boundary. We found that in collisions where plasmid-carrying individuals (purple) are highly intermixed with plasmid-free (green) individuals in the donor biofilm, most of the plasmid-free cells (green) become plasmid-carrying cells (cyan) due to intraspecific transfer, increasing the maximal plasmid load at the collision boundary (Fig. 4B). We also found that, for the highly intermixed scenario, both the total number of new transconjugants (blue) and the number of plasmid transfer events at the collision boundary were higher than in the low-intermixed scenario (two-sample two-sided Welch test; *P* = 0.021, *P* = 0.007, n = 5) (Fig. 4D). Next, we tested the effects of intermixing of purple and green cells on the intermixing of new transconjugants (blue) and recipient cells (grey) in the recipient biofilm, and further effects on plasmid spread within the recipient biofilm (Fig. 4C). We again found that in collisions where plasmid donors (purple) are highly intermixed with plasmid-free (green) individuals, both the total number of new transconjugants (blue) and the number of plasmid transfer events at the collision boundary are higher (two-sample two-sided Welch test; *P* = 0.0005, *P* = 0.001, n = 5) (Fig. 4E). We then quantified the number of plasmid transfer events that occurred within the recipient biofilm and found that there were significantly more transfer events for highly spatially intermixed conditions (138 ± 21) compared to poorly spatially intermixed conditions (91 ± 28) (two-sample two-sided Welch test; *P* = 0.018, n = 5). Indeed, over one-third of plasmid spread within the recipient colony was due to plasmid transfer events between cells of the same strain (33.7 ± 5.1%).

## Discussion

Linking the determinants of plasmid transfer within and between spatially structured microbial communities is of paramount interest for predicting the spread of antibiotic resistance (AR) and other plasmid-encoded traits in sparse biofilms. In this study, we showed that plasmid-encoded AR can readily transfer between spatially separate biofilms upon their expansion and physical collision (Fig. 1C). The new plasmid-carrying genotype is formed immediately after collision, with the potential to further transfer the plasmid into adjacent plasmid-free regions. We revealed that the initial distance between expanding biofilms and the plasmid load dynamics during biofilm expansion determine the spread of the AR-encoding plasmid into the recipient biofilm (Figs. 2 and 3). Our findings also highlight spatial intermixing between plasmid-carrying and plasmid-free individuals as a key driver of plasmid-mediated AR spread between biofilms (Fig. 4).

In the absence of antibiotic selection, there is a period during which plasmid loss upon cell division and selection for plasmid-free cells decreases the opportunities for plasmid transfer into adjacent biofilms. This is evident as a decaying relationship between the extent of new transconjugants in the recipient biofilm and the distance between biofilm inocula (Fig. 2B). The segregation control system of a particular plasmid will have a large impact on its temporal persistence in a given biofilm [39, 40], and the relative cost of the plasmid will also determine how rapidly plasmid-free cells dominate the expansion frontier due to increased relative fitness [41]. There are many cases in which plasmids are able to persist within the host strain even in the absence of a positive selection pressure due to compensatory mutations that eliminate the costs of plasmid carriage [42, 43]. In such situations we expect transfer to be independent of the initial distance between biofilms provided these enter in physical contact. This means that the taxonomic composition of the biofilm and the biology of the AR-encoding plasmids will have a large impact on plasmid spread between adjacent biofilms [44].

We found that the spatial intermixing of plasmid-carrying and plasmid-free cells at the expansion frontier of a donor biofilm affects both the total plasmid load and the degree of intermixing of the plasmid-free and plasmid-carrying individuals in the recipient biofilm. More intermixed populations lead to a larger number of cell-cell contacts between phenotypically distinct types (in this case antibiotic resistant and sensitive individuals). For a contact-dependent process such as plasmid conjugation, this leads to a larger number of possible non-redundant transfer events (i.e., transfer events from plasmid donor to potential recipient cells as opposed to transfer events from plasmid donor to adjacent plasmid donor cells), which results in a larger plasmid spread [18]. This explains why we observed a linear increase in the number of new transconjugants after biofilm collision (Fig. 3D), but a plateau in the number of new transconjugants being formed at the collision boundary (Fig. 3E), because after all potential donor-recipient cell contacts at the collision boundary were realized there were no more targets for additional plasmid transfer.

In the light of our findings, in natural systems such as the human gut, processes leading to higher spatial intermixing between populations are expected to increase plasmid spread. For example, in the mouse gut, the spread of AR-encoding plasmids is maximized with the frequent contact of persisting plasmid donors with invading plasmid-free enteric pathogens [45, 46]. Frequent physical disturbances increase the number of new donor-recipient contacts by reshuffling the spatial positioning of cells and promote widespread plasmid transfer [15]. Upon physical contact between biofilms, however, we confirmed both experimentally and theoretically previous work that suggested transfer only occurs at biofilm boundaries [11], as shown by the narrow extent of the new *E. coli* transconjugants at the collision boundary. Seoane et al. [47] reported similar results, where there was plasmid transfer between small cell colonies, but plasmid invasion was limited by the inactivity of cells and the physical compression towards the colony center. This localized transfer can still be very relevant for the maintenance of unstable plasmids in habitats where disturbances or environmental gradients promote the dispersal and regrowth of spatially structured biofilms associated for example to the mammalian gut, plant structures, or surfaces in aquatic environments (e.g. for gradients in oxygen concentrations; [48]).

This link between intra- and inter-biofilm dynamics sets a basis for understanding antibiotic resistance spread at the metacommunity level, which better resembles processes occurring in natural systems such as sparse biofilms (Fig. 5). The persistence of AR in microbial communities even in the absence of antibiotic pressure could be explained by spatial factors such as the spatial intermixing between cells that operate both within and between biofilms. In spatially structured systems such as the gut lumen or the dental plaque, this conceptualization might be of interest because the spread of plasmid-encoded AR could be modelled based on pre-existing models from metacommunity theory [49, 50]. Source-sink dynamics are a way by which a stable plasmid donor ensures the plasmid is maintained in unstable plasmid recipients by frequent HGT [51]. Our study suggests that depending on the physical proximity between biofilm sources and sinks of plasmid transfer, frequent collisions can lead to the maintenance of plasmids via HGT even when the plasmid is unstable in all community members [51, 52]. This implies that in a metacommunity context the entire metacommunity will have access to the plasmid under source-sink dynamics provided there is frequent physical contact between biofilms.

**Figure 5.**
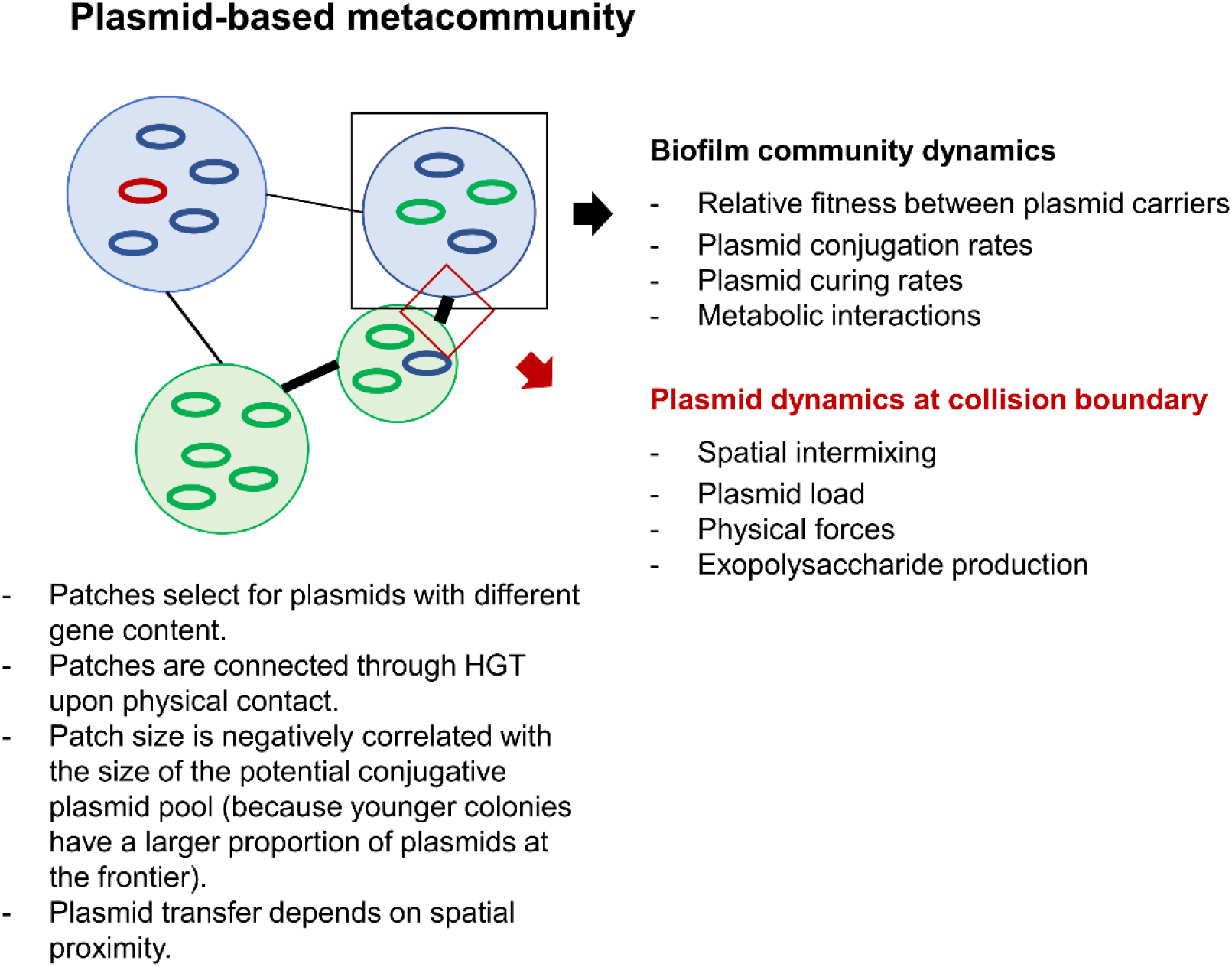
General framework of plasmid spread across a biofilm metacommunity. The scheme summarizes the processes driving the spread of plasmids in spatially structured microbial communities. The width of the lines connecting the communities indicates the magnitude of dispersal of genotypes or plasmids between them. The graph separates the mechanisms that drive plasmid transfer within the communities and at the collision boundaries between them. The spread of plasmid-encoded AR or other traits depends on the load of antibiotic resistance genes and spatial intermixing at the expansion frontier of the colliding biofilms. These respond to multiple processes intrinsic to the dynamics within each of the communities. However, at the collision boundary other forces play an important role because these determine whether the potential horizontal gene transfer can physically occur.

The relative simplicity of our experimental system and modelling approach also has several limitations. Our simplified consortia might not reflect the complex interactions and relative growth differences within complex biofilms. However, the processes we describe here should hold, as different genotypes will persist at the expansion frontier and be susceptible to plasmid transfer. Second, we have not tested the dependency of plasmid transfer on the community composition of the adjacent colliding biofilms. Some taxa are known to produce a thick layer of exopolysaccharides that can prevent cell-cell contacts even when there are compressive physical forces at play [53]. Also, some taxa might not have compatible conjugation machineries to establish pili junctions required for effective transfer [54], while some biofilms will prevent collisions in the first place via chemical signaling [55]. The generalizability of our findings could be confirmed with studies that implement our simplified experimental system using diverse sets of strains from multiple taxonomic groups and traits. There are also multiple factors we could not address that will determine the extent of this spread into the recipient biofilm. First, physical forces might push new transconjugants towards the collision boundary, uplifting them and creating a vertical rather than a horizontal expansion [49]. Second, the metabolic state of cells closer to the center of the biofilm will also determine their ability to capture and transfer the plasmid. Here nutrient availability, interspecies interactions, and abiotic stressors might play an important role at determining such cellular activity. Third, the presence of even subinhibitory concentrations of the antibiotic, to which a plasmid confers resistance, can create a positive selective pressure that promotes further transfer.

We believe our results provide proof of principle and support for an expanded framework of plasmid-mediated antibiotic resistance spread in spatially structured microbial landscapes. Future research should test the influence of subinhibitory antibiotic pressure and nutrient availability in experimental systems using more taxonomically diverse biofilms. Furthermore, *in vivo* imaging of actual biofilm collisions would provide quantitative information about the frequency and extent of antibiotic resistance spread via plasmid conjugation between adjacent biofilms under clinically and environmentally relevant conditions.

## Materials and methods

### Bacterial strains and plasmid

The initial consortium carrying antibiotic resistance consisted of two genetically engineered mutants of the bacterium *Pseudomonas stutzeri* A1501, whose genetic modifications and growth traits have been previously described [56, 57]. Each strain is genetically identical to the other except for containing a different isopropyl β-D-1-thiogalactopyranoside (IPGT)-inducible fluorescent protein-encoding gene located on the chromosome (*egfp* encoding for green fluorescent protein or *echerry* encoding for red fluorescent protein (Supplementary Table 1), which enables us to distinguish and quantify the different strains when grown together. The methods used to construct the strains have been described in detail elsewhere [57]. The strain carrying the *echerry* gene was initially transformed with the R388-derivative plasmid pAR145 (pSU2007 *aph*::*cat*-P_A1/04/03_-*cfp*\*-T_0_, described in [58]). Plasmid pAR145 encodes for chloramphenicol resistance and is marked with an IPTG-inducible *ecfp* gene (encoding for cyan fluorescent protein). The potential recipient strain was *E. coli* DH5α [F2 *supE44 lacU169* (*w80lacZDM15*) *hsdR17 recA1 endA1 gyrA96thi-1 r elA1*] [59].

### Biofilm collision experiments

We performed biofilm collision experiments where we inoculated a mixture of the *P. stutzeri* strains and the *E. coli* DH5α strain as single droplets onto agar plates and allowed them to grow into colonies (biofilms) until collision. We initially grew all the strains separately in oxic liquid lysogeny broth (LB) medium overnight at 37°C and equalized their optical densities at 600nm (OD_600_) to 2. We next mixed the two *P. stutzeri* strains to a fixed 1:1 ratio (vol:vol). The *echerry*-marked donor strain contained pAR145, which encodes for *ecfp,* and thus displayed the composite color purple (red and blue). The *egfp*-marked potential recipient strain only displayed the color green. Using a Tecan Evo 200 liquid handling system (Tecan, Männedorf, Zurich, Switzerland), we then deposited pairs of droplets of the *P. stutzeri* mixture and the *E. coli* strain (1 μl of each culture) at four discrete spatial positions on each LB agar plate adjusted to a pH of 7.5 and amended with 1 mM IPTG. We programmed the liquid handling system to deposit the *P. stutzeri* mixture and *E. coli* droplets at distances between droplet centroids of 2.80, 2.90, 3.00, 3.10, 3.20, 3.30, 3.40, 3.50, 3.70, and 4.00 mm with 3 replicates per distance. We finally incubated the plates for 96h under room temperature in oxic conditions.

### Bacterial colony imaging and quantitative image analysis

We imaged the colonies immediately upon completion of the incubation period using a Leica TC5 SP5 II confocal microscope (Leica Microsystems, Wetzlar, Germany) with objectives 10x/0.3na (dry) and 63 x/1.4na (oil) (Etzlar, Germany). We scanned the entire biofilms (*P. stutzeri* mixture and *E. coli*) by stitching together multiple frames of 1024 x 1024 pixels. We set the laser emissions to 514 nm for the excitation of the red fluorescent protein, 488 nm for the excitation of the green fluorescent protein, and 458 nm for the excitation of the cyan fluorescent protein. We analyzed the images in ImageJ https://imagej.nih.gov/ij/) using FIJI plugins (v. 2.1.0/1.53c; https://fiji.sc).

We quantified plasmid dosage (i.e., the extent cyan fluorescent protein-expressing cells) using the “Sholl analysis” plugin [60] on the binarized image of the blue channel after application of a noise-reduction threshold of 20 pixels. The “Sholl analysis” calculated the number of blue pixels at 10 μm radial increments from the centroid to the colony edge. We initiated the analysis at 1500 microns from the colony centroid because fluorescent signals at smaller radii could not be precisely resolved by image analysis. We then applied the “Area to line” function of the “Overlay” plugin to register the number of blue pixels (here coded as 255 because pixel intensity was not the main target) and background (coded as 0) for the areas corresponding to each 10 μm radial increment. We defined plasmid dosage as the total number of blue pixels divided by the total number of pixels along each radius. We performed all downstream analyses in R Studio (v1.3.1073, https://www.rstudio.com).

### Individual-based modelling of antibiotic resistance spread between microbial communities on surfaces

We performed individual-based computational modelling of microbial range expansions using CellModeller 4.3, which is a computational framework designed to physically and chemically simulate rod-shaped cells with user-defined rules [38]. We modelled individual rod-shaped cells as three-dimensional capsules (i.e., cylinders with hemispherical ends), where capsules grow by extending their length and experience frictional drag that stops them from growing into one another. We used the default setting for the parameter that controls frictional drag (gamma = 10). As cells grow, they add a constant volume until reaching a critical size defined by default settings where they then divide into two daughter cells. Meanwhile, cells have limited potential of shoving due to physical forces that limit the displacement of cells upon division. In CellModeller, cells are abstracted as computational objects referred to as a cellState (cs) that contain all the information regarding an individual cell, including its spatial position (pos[x, y, z]), rotational orientation (dir[x, y, z]), cell length (len), growth rate (growthRate), and cell type (cellType). The cell-type is an arbitrary label that allows us to simulate different cellular behaviors.

In CellModeller, individual cells are modelled as cylinders of length *l* capped with hemispheres that result in a capsule shape, with both hemispheres and the cylinder having a radius *r.* At each simulation step, a cell increases in length based on its growth rate parameter, which is physically constrained by the other cells in its physical proximity. In this work, we initiated cells to have *r* = 0.04 and *l* = 2 and set cells to divide when their length reaches the critical division length *l_div_* with the following equation:

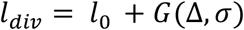

where *l*_0_ is the initial cell length at birth and *G* is a random gaussian distribution with mean Δ = 2 and standard deviation *σ* = 0.45. Therefore, when a cell divides, the two daughter cells are initiated with *l_div_*/2 and a new target division length is assigned to each daughter cell calculated from the equation above. The addition of constant mass has been found to accurately model bacterial division while maintaining cell size homeostasis as described elsewhere [61].

We extended our model to incorporate plasmid transfer and plasmid loss. As part of the biophysics in CellModeller, physical contacts between cells are recorded at each step to minimize any overlap between cells. We altered the code such that each cell kept track of their contacts, which allowed us to model plasmid transfer when cells were in contact. We activated this function by setting the argument ‘compNeighbours=True’ when initiating the biophysical model. For the simulations in Fig. 3, we initially positioned three cell types – green plasmid-free cells (cellType 0), purple plasmid-carrying cells (cellType 1) and grey plasmid-free cells (cellType 2) by loading 394 cells in total (101 green cells, 96 purple cells, and 197 uncolored cells) across the grid with a uniform distance of 5 units between cells along the x and y axes. We loaded green, purple, and uncolored cells according to the checkerboard arrangement within a circle of radius 40 units. Initially, we set the origin of one “colony” at coordinate (−25, −25) and the other at (25, 25). In order to allow two “colonies” to collide at a later time, we adjusted the distance between the two by increasing the absolute value of the x- and y-axes to (±30, ±30), (±35, ±35) and (±40, ±40). We loaded all the cells with the z coordinate set to 0. Thus, we constrained their orientations and dynamics to the x, y plane. We tested different plasmid transfer and loss probabilities and selected a plasmid loss probability P_l_ = 0.0005 and a plasmid conjugation probability P_c_ = 0.005 to capture important features from the experimental results (Fig. 2A). We set the growth rate of green cells, purple cells and grey cells to 1, 0.83 and 0.8, respectively, such that the relative growth rates were consistent with our experimental measurements. For the simulations in Fig. 4, we kept most of the primary settings from those used for the simulations in Fig. 3 but we removed the plasmid loss process. We kept all cellTypes growing at the same rate 1) because the length and time scales are small when simulating collision boundaries, and 2) to isolate the effects of spatial positioning from differential growth rates.

### Quantification of pAR145 transfer and loss rates

We estimated pAR145 transfer (conjugation) rates between all experimental strains by growing three pairs of pAR145 donor and potential recipient strains (purple with green, purple with recipient *E. coli*, and transconjugant *E. coli* with recipient *E. coli*) on filters with a pore size of 0.22 μm (Merck Millipore) for 24 hours at room temperature. We started the donor and recipient cultures independently and then adjusted each culture to OD_600_ = 2 using 0.89% (w/v) sodium chloride solution and mixed them at a ratio of 1:1 v/v. We then evenly spread the 50 μL mixtures onto filters applied directly to the surfaces of agar plates, where the filters increase physical contacts between cells. After 24 hours of incubation at room temperature, we washed off the cells from the filters using phosphate-buffered saline (PBS). We then quantified the number of new transconjugants by serially diluting the PBS solution, spreading it onto LB agar plates supplemented with 25 μg mL^-1^ chloramphenicol and the corresponding selective antibiotic (50 μg mL^-1^ gentamycin for *P. stutzeri,* 25 μg mL^-1^ nalidixic acid for *E. coli* DH5α), and incubating the plates at 37°C for 24h. We then counted the number of chloramphenicol resistant colonies and estimated transfer rates as the ratio of new transconjugants to the total number of pAR145 carriers and pAR145-free individuals as described in [62].

We estimated pAR145 loss probabilities by growing pAR145-carrying strains in a shaking 96-well plate reader using an Eon™ Microplate Spectrophotometer with high performance microplate-based absorbance readings. We followed the methods described in [63] to screen for colonies that lost pAR145 after three hours of growth at 37 °C. We kept the growth duration short because plasmid loss probabilities are commonly measured by quantifying the accumulation of plasmid-free cells over time, which consists of both plasmid loss events and the subsequent growth of plasmid-free cells. Therefore, we minimized subsequent growth by shortening the incubation time. Next, we spread 50 μL of a pAR145-carrying culture that was grown and adjusted as described above for estimating pAR145 transfer probabilities. After 24h of incubation at 37°C, we counted the number of colonies with and without pAR145 and estimated the pAR145 loss probabilities as the ratio of pAR145-free to pAR145-carrying individuals.

### Quantification of relative growth rates between pAR145-carrying and -free strains

We estimated the relative growth rates between pAR145-carrying and -free strains using a colony collision method bases on colony geometry described in [64]. We used the pipetting robot to place two 1 μL droplets (one droplet for each strain tested) 3 mm apart from each other on an LB agar plate, where the cell density of each droplet was set to OD_600_ = 2. We next incubated the LB agar plates under room temperature for 96h to allow the droplets to form colonies and for the colonies to collide into each other. We then used the geometric approach described in [64] to estimate the relative growth rates of the strains based on the arc of the collision boundary between the two corresponding colonies. Briefly, the formula we used is:

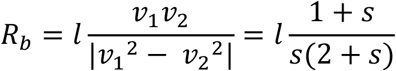

where *l* is the distance between two colonies; s is the selective advantage or cost that pAR145 confers; *v*_1_ is the expansion velocity of the pAR145-free colony (faster velocity); *v*_2_ is the expansion velocity of the pAR145-carrying colony (slower velocity); *R_b_* is the radius of the circle generated by the arc at the boundary. Measurements of *R_b_* and *l* are sufficient to derive s. We quantified *R_b_* and *l* for 4 replicates using Adobe Illustrator 2022 where we manually drew lines and circles to extract the values. While the image scale can differ among replicates, *R_b_* and *l* are proportional in one image i and s will therefore not be affected.

## Acknowledgments

We thank Dr. María Pilar Garcillán-Barcia (University of Cantabria) for providing I plasmid pAR145 and *E. coli*DH5α. J.R. acknowledges funding from the Swiss National Science Foundation (Early Postdoc Mobility grant: P2EZP3_199849). M.M. acknowledges funding from the Swiss National Science Foundation (Ambizione grant: 1 PZ00P3_180147).

## Supplementary materials

**Supplementary Table 1.**
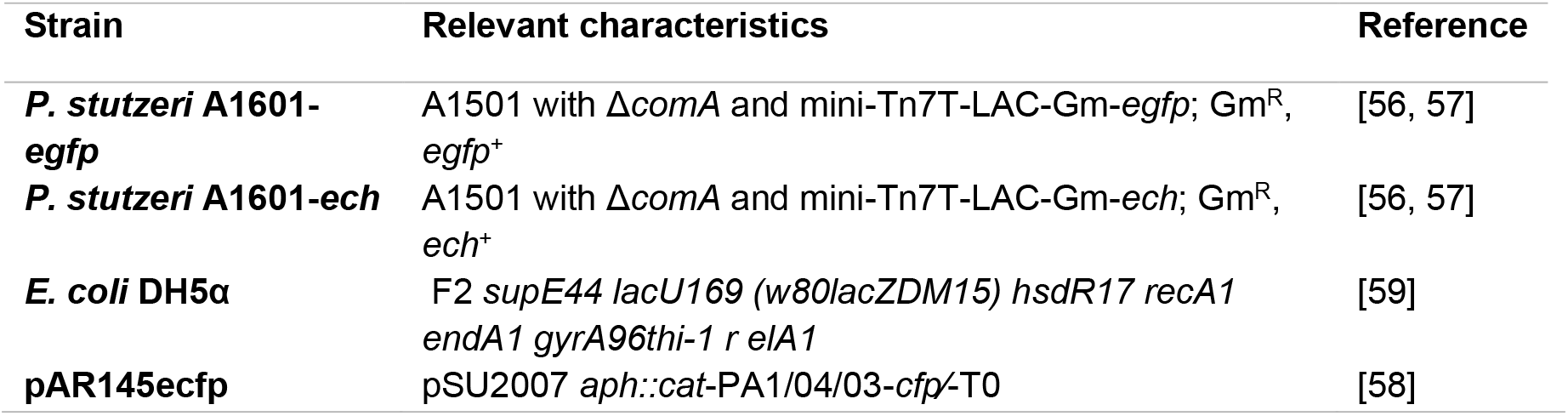
Specifications of the strains and plasmid used in this study.

